# Effects of mucin and DNA concentrations in airway mucus on *Pseudomonas aeruginosa* biofilm recalcitrance

**DOI:** 10.1101/2022.03.24.485733

**Authors:** Kaitlyn R. Rouillard, William J. Kissner, Matthew R. Markovetz, David B. Hill

## Abstract

The pathological properties of airway mucus in cystic fibrosis (CF) are dictated by mucus concentration and composition, with mucins and DNA being responsible for mucus viscoelastic properties. As CF pulmonary disease progresses, the concentrations of mucins and DNA increase and are associated with increased mucus viscoelasticity and decreased transport. Similarly, the biophysical properties of bacterial biofilms are heavily influenced by the composition of their extracellular polymeric substances (EPS). While the roles of polymer concentration and composition in mucus and biofilm mechanical properties have been evaluated independently, the relationship between mucus concentration and composition and the biophysical properties of biofilms grown therein remains unknown. *Pseudomonas aeruginosa* biofilms were grown in airway mucus as a function of overall concentration and DNA concentration to mimic healthy and CF physiology and biophysical properties were evaluated with macro- and microrheology. Biofilms were also characterized after exposure to DNase or DTT to examine the effects of DNA and mucin degradation, respectively. Identifying critical targets in biofilms for disrupting mechanical stability in highly concentrated mucus may lead to the development of efficacious biofilm therapies and ultimately improve CF patient outcomes. Overall mucus concentration was the predominant contributor to biofilm viscoelasticity and both DNA degradation and mucin reduction resulted in compromised biofilm mechanical strength.

## Introduction

Chronic respiratory infection by *Pseudomonas aeruginosa* is a major contributor to morbidity and mortality in cystic fibrosis (CF) (1–3). The CF airway is characterized by abnormal mucus properties including hyperconcentration, increased viscoelasticity, and decreased mucociliary transport (4–6). Mucins, high molecular weight glycoproteins, form a polymeric mesh network that is responsible for the viscoelastic properties of mucus (4, 7). Abnormal water absorption, mucus dehydration, and mucin hypersecretion in CF together result in highly concentrated mucus in the airway which promotes chronic respiratory infections (5, 8). Mucus in CF represents an ideal growth environment for *P. aeruginosa* as it both provides a physical mesh to grow within and acts as a nutrient source (5, 9). High density gels such as >0.6% agarose and CF sputum promote bacterial aggregation (8), and attachment to mucins has been hypothesized to improve biofilm stability (5, 8, 10). Indeed, pathological mucus properties are associated with increased incidence of infection, and *P. aeruginosa* in particular establishes biofilms, which are viscoelastic aggregates of bacteria protected by a matrix of extracellular polymeric substances (EPS) (5, 11, 12). The composition of biofilm EPS is dependent upon a number of factors including the biophysical and biochemical properties of the growth environment (13–15).

Both mucus and biofilms have viscoelastic properties that are heavily influenced by the concentration and composition of polymeric substances (4, 11). For mucus, those polymers include DNA, which is released into the airway as a result of the chronic inflammatory response, and mucins (16, 17). In biofilms, the EPS include DNA that bacteria self-produce or uptake from the environment, polysaccharides, lipids, and proteins (13). The importance of DNA in both mucus and biofilm biophysical properties has recently been investigated and correlated with decreased transport of mucus (17) and antibiotic susceptibility in biofilms (2, 18). Environmental DNA has been associated with increased biofilm viscoelasticity and decreased susceptibility to mechanical and chemical challenge (2, 18, 19). However, the role of DNA in airway mucus in influencing biofilm biophysical properties has yet to be elucidated.

In general, the relationship between airway mucus composition and biofilm recalcitrance is poorly understood. It has been established independently that increased mucus concentration promotes biofilm formation (5, 8) and that increased DNA concentrations are associated with more robust biofilms (2, 18, 19), but biofilm mechanical properties as a function of DNA concentrations in mucus have not yet been evaluated. It is essential to understand how changes in mucus composition and concentration affect biofilm mechanics and susceptibility in order to identify potential therapeutic targets and improve patient outcomes. Conventional antibiotics are limited by poor diffusion through and reduced efficacy in mucus (20, 21). Additionally, the rise in antibiotic resistance limits the spectrum of antibiotics available to use for chronic infections, particularly as biofilms inherently exhibit decreased susceptibility to antibiotics compared to their planktonic counterparts (22, 23). Improving our understanding of biofilm mechanics will ultimately lead to more effective treatment design.

Herein, we describe the evaluation of *P. aeruginosa* biofilm mechanical strength with macro- and microrheological techniques as a function of human bronchial epithelial (HBE) mucus composition. Mucus at 2% and 5% solids to represent healthy and pathological mucus, respectively, was supplemented with salmon sperm DNA at 100:1 and 20:1 mucin to DNA ratios to mimic early and more advanced CF mucus (17, 24). Biofilms were also exposed to the DNA degradation enzyme, DNase, or to a reducing agent, dithiothreitol (DTT), to investigate: 1) the relative contribution of DNA to biofilm mechanical strength, and 2) the disruption of chemical bonds and physical interactions in the biofilm matrix. Identifying key targets for biofilm disruption as a function of mucus composition may improve the selection and efficacy of future antibiofilm treatments in CF pulmonary disease.

## Results

### The combination of mucins and DNA produce mechanically strong biofilms in nutrient broth

To first evaluate the individual contributions of mucins and DNA to biofilm mechanical strength, *P. aeruginosa* biofilms were grown in TSB supplemented with porcine gastric mucins (PGM) and/or salmon sperm DNA at concentrations used for making artificial sputum media (ASM) (25). Combinations of PGM and DNA were made with PGM at 5 mg/mL and mucin to DNA ratios of 100:1 (i.e., 0.05 mg/mL DNA) and 20:1 (i.e., 0.25 mg/mL) to mimic mucin to DNA ratios observed in CF in preschool aged children and adults, respectively (17). *Pseudomonas aeruginosa* biofilms exhibited viscoelasticity with an elastic modulus (G’) greater than the viscous modulus (G”), meaning they behave as a soft viscoelastic solid, i.e., tanδ <0.5 (26). Biofilms grown in TSB with salmon sperm DNA were statistically indistinguishable at either DNA concentration (Table 1). However, when grown in TSB with PGM, biofilms exhibited decreased complex viscosity (η*, 0.05 Pa·s) compared to TSB alone (0.10 Pa·s). A similar phenomenon has been described previously wherein low concentrations of mucins have been shown to prevent biofilm formation (27–29). In contrast, the combination of PGM and DNA resulted in mechanically stronger biofilms compared to TSB alone, with η* values being 3-4x greater. Both mucin to DNA ratios of 100:1 and 20:1 produced biofilms with statistically similar η* values, 0.31 and 0.41 Pa·s, respectively, though the larger DNA concentration (20:1) was associated with a greater absolute value of η*.

**Table 1.** Macroscopic moduli of *P. aeruginosa* biofilms grown in TSB as a function of mucin and DNA concentrations. Data is presented as the mean and standard deviation of ≥3 separately grown and analyzed biofilms.

| Growth Media | G' (Pa) | G'' (Pa) | $\eta^*$ (Pa·s) |
| --- | --- | --- | --- |
| TSB | $0.64 \pm 0.27$ | $0.11 \pm 0.03$ | $0.10 \pm 0.04$ |
| DNA 0.5 mg/mL | $0.78 \pm 0.43$ | $0.18 \pm 0.08^*$ | $0.13 \pm 0.07$ |
| DNA 5 mg/mL | $0.78 \pm 0.52$ | $0.18 \pm 0.12$ | $0.14 \pm 0.09$ |
| PGM 0.5 mg/mL | $0.21 \pm 0.11^*$ | $0.06 \pm 0.02$ | $0.04 \pm 0.02^*$ |
| PGM 5 mg/mL | $0.33 \pm 0.15^*$ | $0.07 \pm 0.03$ | $0.05 \pm 0.02^*$ |
| $\nabla$PGM:DNA 100:1 | $1.86 \pm 1.40^*$ | $0.47 \pm 0.32^*$ | $0.31 \pm 0.23^*$ |
| $\nabla$PGM:DNA 20:1 | $2.50 \pm 0.49^*$ | $0.71 \pm 0.34^*$ | $0.41 \pm 0.25^*$ |

\* $p < 0.05$ compared to TSB biofilms
$\nabla$ PGM held at 5 mg/mL and DNA adjusted to ratio

To elucidate the importance of native DNA in biofilm mechanical stability, TSB biofilms were exposed to the DNA degradation enzyme, DNase, and evaluated with particle tracking microrheology (PTMR). Microrheology is capable of resolving spatial heterogeneity that is lost in the bulk measurement of macrorheology (30). The degradation of native DNA within the biofilm was hypothesized to compromise structural integrity. Exposing TSB biofilms to DNase resulted in significant changes in biofilm microrheology (Figure 1), including mean η* and heterogeneity, as demonstrated by the swarm charts in Figure 1A. Quantification of heterogeneity was performed using the non-Gaussian parameter, κ, wherein greater κ values indicate more heterogeneity within the sample (31). For normal Brownian motion, the value of κ should be ≪1 (31). Values of κ for each DNase concentration are listed in Figure 1A. The most homogeneous sample was biofilms treated with 100 μg/mL DNase, though all doses of DNase were associated with decreased κ values. The lowest tested dose, 10 μg/mL, had no significant effect on biofilm complex viscosity, η* (Figure 1B), but 20 μg/mL increased η*. The greatest doses of DNase, 100 and 150 μg/mL, resulted in η* values greater than that of the PBS-treated control but lower than that for 20 μg/mL DNase. No tested concentration of DNase reduced biofilm η* to that of water (1 mPa·s), indicated by the dashed line in Figure 1C. Thus, greater concentrations of DNase may be needed to fully dissociate the biofilm.

**Figure 1.**
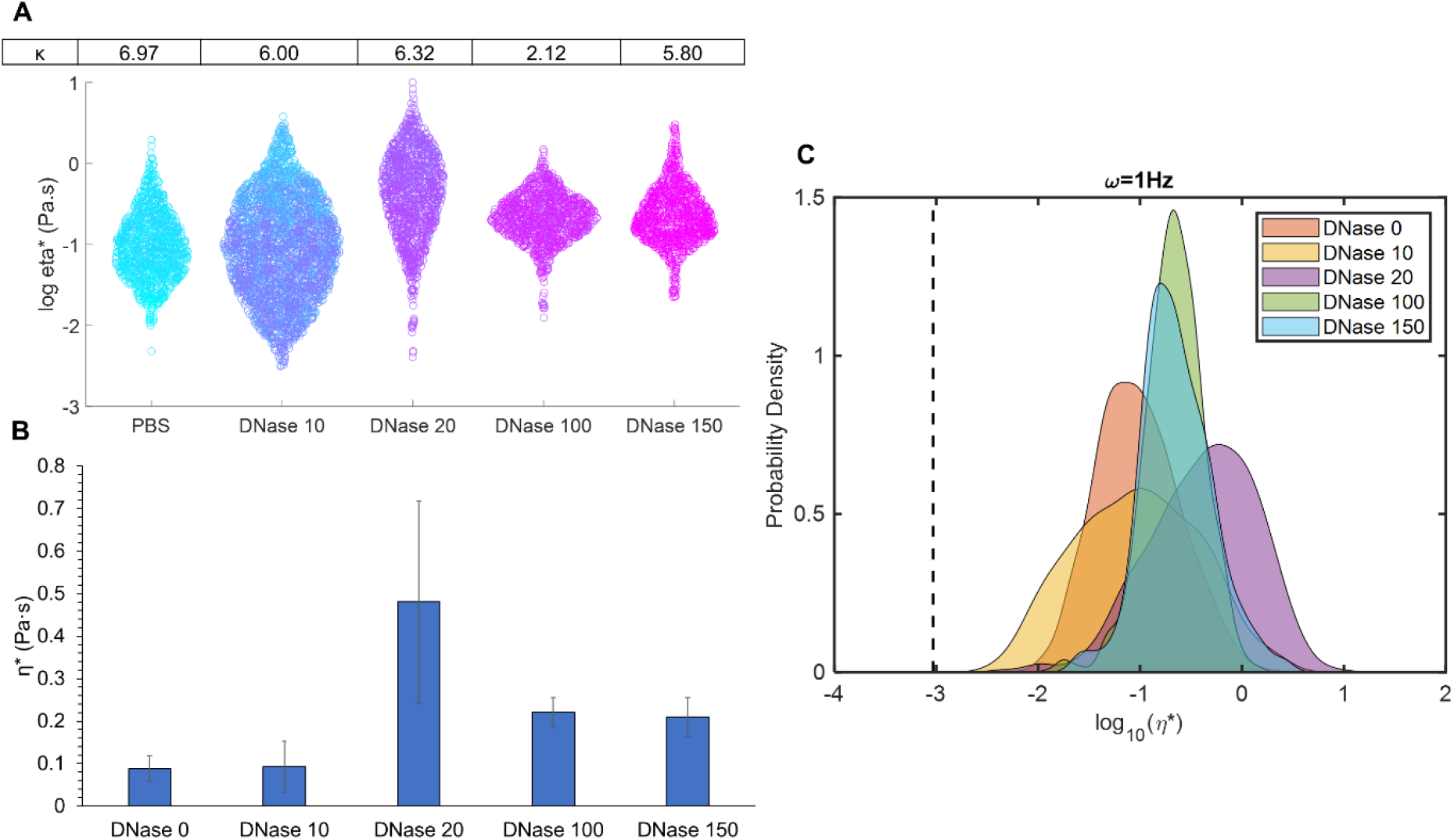
Biofilms grown in TSB microrheology as a function of DNase concentration. (A) Swarm chart of all tracked bead η* measurements in treated biofilms with κ values listen in table above corresponding dose of DNase. (B) Mean ensemble η* and standard deviation of treated biofilms. (C) Probability distribution of each bead η* for each condition. The viscosity of water is indicated by the dashed line. Data is presented for all beads tracked in ≥3 separately grown and analyzed biofilms.

### Increasing overall mucus concentration and DNA to mucin ratios resulted in increased viscoelasticity

To more accurately mimic CF physiology, mucus collected from human bronchial epithelial (HBE) cultures was selected as a biofilm growth media. First, the biophysical properties of mucus were evaluated as a function of salmon sperm DNA concentrations. A comparison of the macro and microrheology of 2% and 5% mucus demonstrated that more concentrated mucus exhibited greater viscoelastic moduli (Table 2), in keeping with previous work (5, 16). On the macro scale, the complex viscosity (η*) of 5% mucus was greater than that of 2% by a factor of ~3. The addition of DNA did not significantly increase η* in 2% mucus but macroscopic moduli were significantly greater in 5% mucus with DNA. Both G’ and G” values trended with η* and were significantly increased in 5% mucus with DNA. Particle tracking microrheology showed that η* was greater in mucus with DNA at both 2% and 5% (Figure 2), though the most significant contributor to η* was overall concentration. In addition, 2% mucus was characterized by a bimodal distribution of lower η*, more water-like component and a higher η*, more solid-like component (Figure 2A), in agreement with previous work (32). This bimodality was likewise observed in 2% mucus with DNA supplementation, suggesting that the addition of DNA at these concentrations is insufficient to homogenize mucus η* values into a single peak that is observed in greater concentrations of HBE mucus (4).

**Table 2.** Mucus macroscopic moduli as a function of concentration and composition. Data is presented as the mean and standard deviation of ≥3 separately prepared and analyzed mucus samples.

| Mucus | $G'$ (Pa) | $G''$ (Pa) | $\eta^*$ (Pa·s) |
| --- | --- | --- | --- |
| 2% HBE | $0.465 \pm 0.200$ | $0.227 \pm 0.039$ | $0.082 \pm 0.031$ |
| 2% 100:1 | $0.523 \pm 0.343$ | $0.195 \pm 0.077$ | $0.090 \pm 0.055$ |
| 2% 20:1 | $0.464 \pm 0.346$ | $0.173 \pm 0.101^*$ | $0.079 \pm 0.057$ |
| 5% HBE | $0.838 \pm 0.354$ | $0.451 \pm 0.074$ | $0.153 \pm 0.055$ |
| 5% 100:1 | $1.782 \pm 0.642^*$ | $1.055 \pm 0.494^*$ | $0.330 \pm 0.128^*$ |
| 5% 20:1 | $1.986 \pm 0.823^*$ | $1.586 \pm 0.174^*$ | $0.418 \pm 0.083^*$ |

\* $p < 0.05$ compared to same concentration mucus without DNA

**Figure 2.**
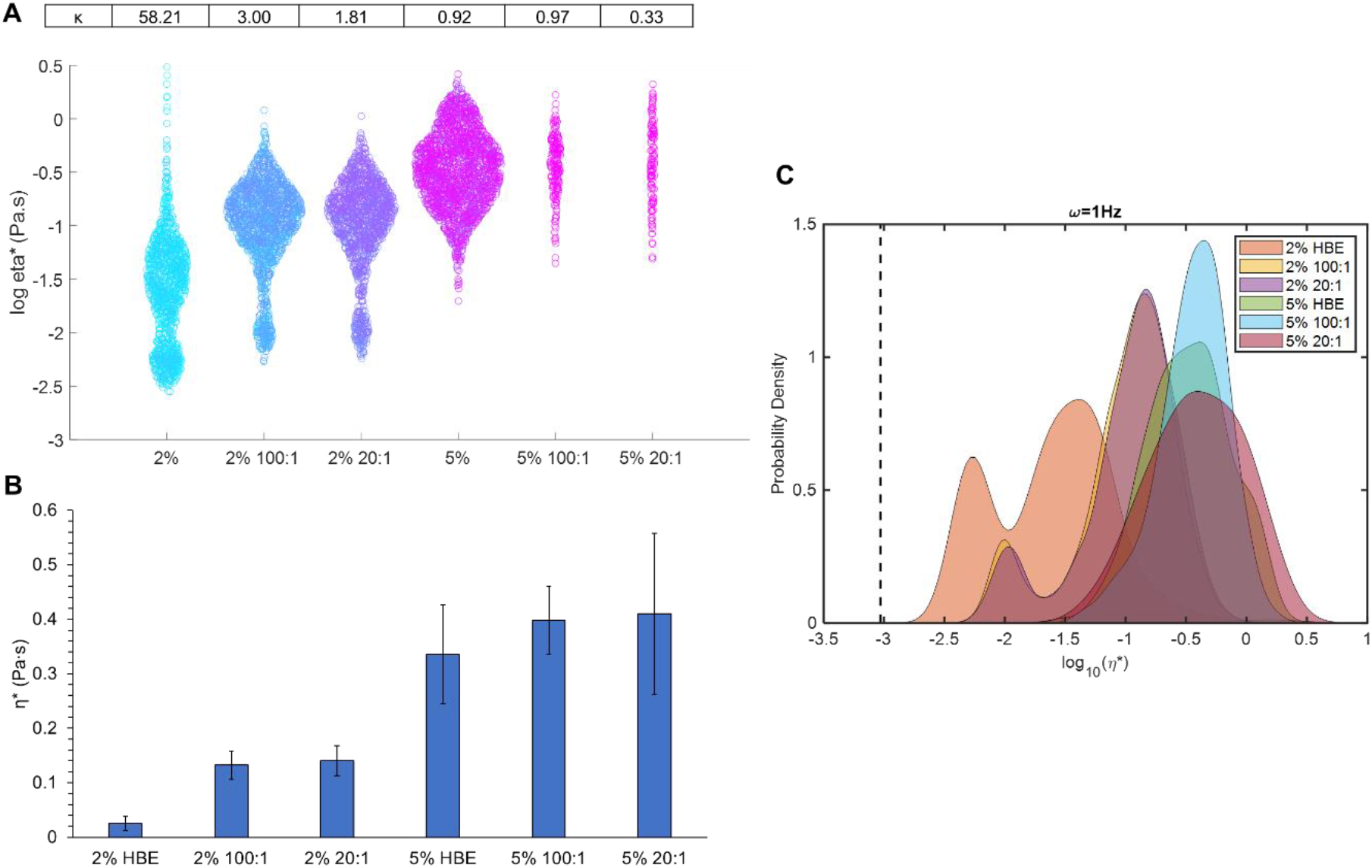
Mucus microrheology as a function of concentration and composition. Swarm chart of all tracked bead η* measurements in each mucus type. The non-Gaussian parameter κ is listed above each mucus type. (B) Mean ensemble η* and standard deviation for each mucus type. (C) Probability distribution of each bead η* for each condition. The viscosity of water is indicated by the dashed line. Data is presented as the distribution of all tracked beads in ≥3 separately prepared and analyzed samples.

### Biofilm rheology is correlated with mucus rheology

After mucus characterization, *P. aeruginosa* biofilms were grown in HBE mucus at 2% and 5% total solids with no DNA, 100:1 mucin to DNA, or 20:1 DNA. Biofilms grown in mucus were rheologically similar to those grown in nutrient broth (Tables 1 and 3). All biofilms were dominated by G’ (tanδ < 0.5), similar to mucus. The presence of DNA in mucus was not associated with any significant changes in macroscopic moduli (Table 3). The predominant discriminator between biofilm rheology was overall mucus concentration, with 5% mucus types facilitating the growth of more robust biofilms (i.e., greater G’, G”, and η* values). The same trend was observed with PTMR wherein biofilms grown in 5% mucus types exhibited one peak at a greater mean η*compared to those grown in 2% mucus types (Figure 3). Consistent with mucus rheology (Figure 2), biofilms grown in 2% mucus types also demonstrated a bimodal distribution into distinct lower and greater η* groupings (Figure 3A). The addition of DNA to 5% mucus was correlated with a greater mean biofilm η*, and biofilms were more heterogeneous compared to those grown in the absence of DNA. Biofilms grown in 5% had statistically greater macroscopic moduli compared to those grown in 2%, but no differences were measured with the addition of DNA at any tested concentration.

**Table 3.**
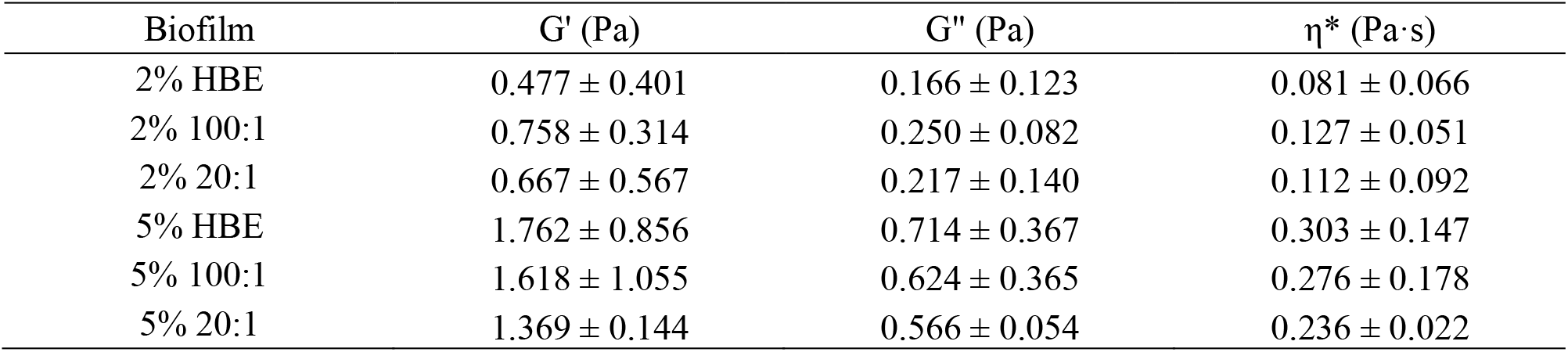
Biofilm macroscopic moduli as a function of mucus concentration and composition. Data is presented as the mean and standard deviation of ≥3 separately grown and analyzed biofilms.

**Figure 3.**
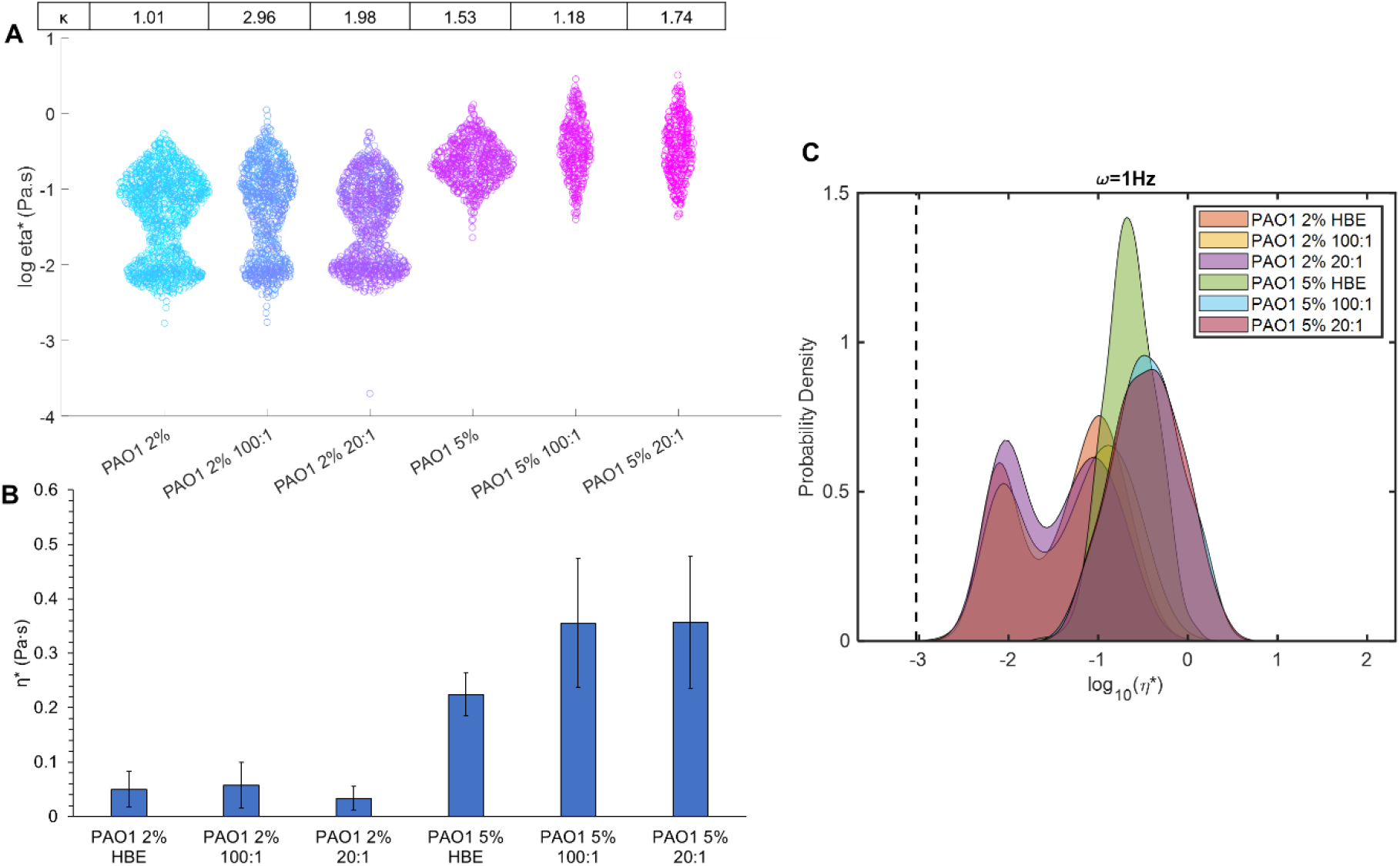
Biofilm microrheology as a function of mucus concentration and composition. (A) Swarm chart of all tracked bead η* measurements in biofilms. The non-Gaussian parameter κ is listed above each biofilm. Dashed line represents the viscosity of water. (B) Mean ensemble η* and standard deviation of biofilms. (C) Probability distribution of each bead η* for each condition. The viscosity of water is indicated by the dashed line. Data is presented for all beads tracked in ≥3 separately grown and analyzed biofilms.

### Treatment with DNase or DTT affects biofilm rheology

Reductions in biofilm macrorheology by DNase or DTT occurred in a predominantly concentration-dependent manner (Table 4). Some concentrations of DNase or DTT caused an increase in biofilm η*, but this phenomenon was not consistent across mucus compositions. For example, 5% HBE biofilm η* significantly increased at 10 μg/mL DNase but was statistically indistinguishable at 100 and 1000 μg/mL. However, DTT reduced 5% HBE biofilm η* at 15 and 150 μg/mL but increased biofilm η* at 1500 μg/mL. Degradation of the biofilm extracellular matrix allows for the disruption and release of interior components such as both live and dead bacteria cells exoenzymes, swellable polysaccharides, proteins, and lipids (13), which can increase bulk rheology. Indeed, biofilm disruption and bacterial cell death via nitric oxide or tobramycin exposure has previously been shown to increase biofilm macrorheology (18). The use of PTMR improves the analysis of local viscosity in the biofilm and describes biofilm heterogeneity (Figures 4 and 5). Biofilms grown in all 2% HBE mucus types were similarly affected by mid (150 μg/mL) to high (1500 μg/mL) doses of DTT (Figure 4). The higher η* component of 2% biofilms disappeared with DTT treatment. Uniquely, the largest dose of DTT reduced 5% biofilm mean η* and was associated with a second peak at lower η* (Figure 5), similar to untreated 2% biofilms. This effect was also observed at the largest dose of DNase.

**Table 4.** Biofilm macroscopic η* after 24 h exposure to DNase or DTT. Data is presented as the mean and standard deviation of ≥3 separately grown and treated biofilms.

| Treatment | 2% HBE | 2% 100:1 | 2% 20:1 | 5% HBE | 5% 100:1 | 5% 20:1 |
| --- | --- | --- | --- | --- | --- | --- |
| PBS | 0.081 ±<br>0.066 | 0.127 ±<br>0.051 | 0.112 ±<br>0.092 | 0.303 ±<br>0.147 | 0.276 ±<br>0.178 | 0.236 ±<br>0.022 |
| DNase<br>10 ug/mL | 0.005 ±<br>0.003 | 0.046 ±<br>0.009 | 0.096 ±<br>0.007 | 1.700 ±<br>0.148 | 0.196 ±<br>0.036 | 0.475 ±<br>0.054 |
| DNase<br>100 ug/mL | 0.009 ±<br>0.004 | 0.052 ±<br>0.009 | 0.018 ±<br>0.004 | 0.250 ±<br>0.024 | 1.243 ±<br>0.116 | 0.591 ±<br>0.045 |
| DNase<br>1000 ug/mL | 0.211 ±<br>0.020 | 0.026 ±<br>0.004 | 0.012 ±<br>0.003 | 0.292 ±<br>0.033 | 0.153 ±<br>0.033 | 0.110 ±<br>0.003 |
| DTT<br>15 ug/mL | 0.030 ±<br>0.004 | 0.130 ±<br>0.081 | 1.337 ±<br>0.629 | 0.105 ±<br>0.003 | 0.122 ±<br>0.009 | 0.292 ±<br>0.012 |
| DTT<br>150 ug/mL | 0.202 ±<br>0.018 | 0.048 ±<br>0.010 | 0.025 ±<br>0.006 | 0.147 ±<br>0.006 | 0.138 ±<br>0.007 | 0.207 ±<br>0.008 |
| DTT<br>1500 ug/mL | 0.477 ±<br>0.068 | 0.049 ±<br>0.012 | 0.059 ±<br>0.036 | 0.477 ±<br>0.068 | 0.085 ±<br>0.002 | 0.158 ±<br>0.004 |

**Table 5.** Biofilm η* measured via PTMR after 24 h exposure to DNase or DTT. Data is presented as the mean and standard deviation of all tracked beads in ≥3 separately grown and treated biofilms.

| Treatment | PAO1 2%<br>HBE | PAO1 2%<br>100:1 | PAO1 2%<br>20:1 | PAO1 5%<br>HBE | PAO1 5%<br>100:1 | PAO1 5%<br>20:1 |
| --- | --- | --- | --- | --- | --- | --- |
| PBS | 0.050 ± 0.033 | 0.058 ± 0.041 | 0.034 ± 0.022 | 0.224 ± 0.039 | 0.355 ± 0.018 | 0.356 ± 0.122 |
| DNase<br>10 $\mu\text{g/mL}$ | 0.026 ± 0.019 | 0.072 ± 0.029 | 0.027 ± 0.020 | 0.153 ± 0.033 | 0.139 ± 0.027 | 0.260 ± 0.064 |
| DNase<br>100 $\mu\text{g/mL}$ | 0.004 ± 0.000 | 0.064 ± 0.023 | 0.025 ± 0.014 | 0.154 ± 0.046 | 0.464 ± 0.175 | 0.199 ± 0.062 |
| DNase<br>1000 $\mu\text{g/mL}$ | 0.004 ± 0.001 | 0.020 ± 0.011 | 0.015 ± 0.010 | 0.232 ± 0.070 | 0.107 ± 0.033 | 0.114 ± 0.033 |
| DTT<br>15 $\mu\text{g/mL}$ | 0.009 ± 0.002 | 0.009 ± 0.001 | 0.028 ± 0.016 | 0.106 ± 0.023 | 0.140 ± 0.036 | 0.199 ± 0.064 |
| DTT<br>150 $\mu\text{g/mL}$ | 0.005 ± 0.000 | 0.007 ± 0.001 | 0.023 ± 0.015 | 0.274 ± 0.091 | 0.134 ± 0.035 | 0.146 ± 0.047 |
| DTT<br>1500 $\mu\text{g/mL}$ | 0.027 ± 0.019 | 0.009 ± 0.002 | 0.007 ± 0.000 | 0.472 ± 0.119 | 0.213 ± 0.105 | 0.132 ± 0.050 |

**Figure 4.**
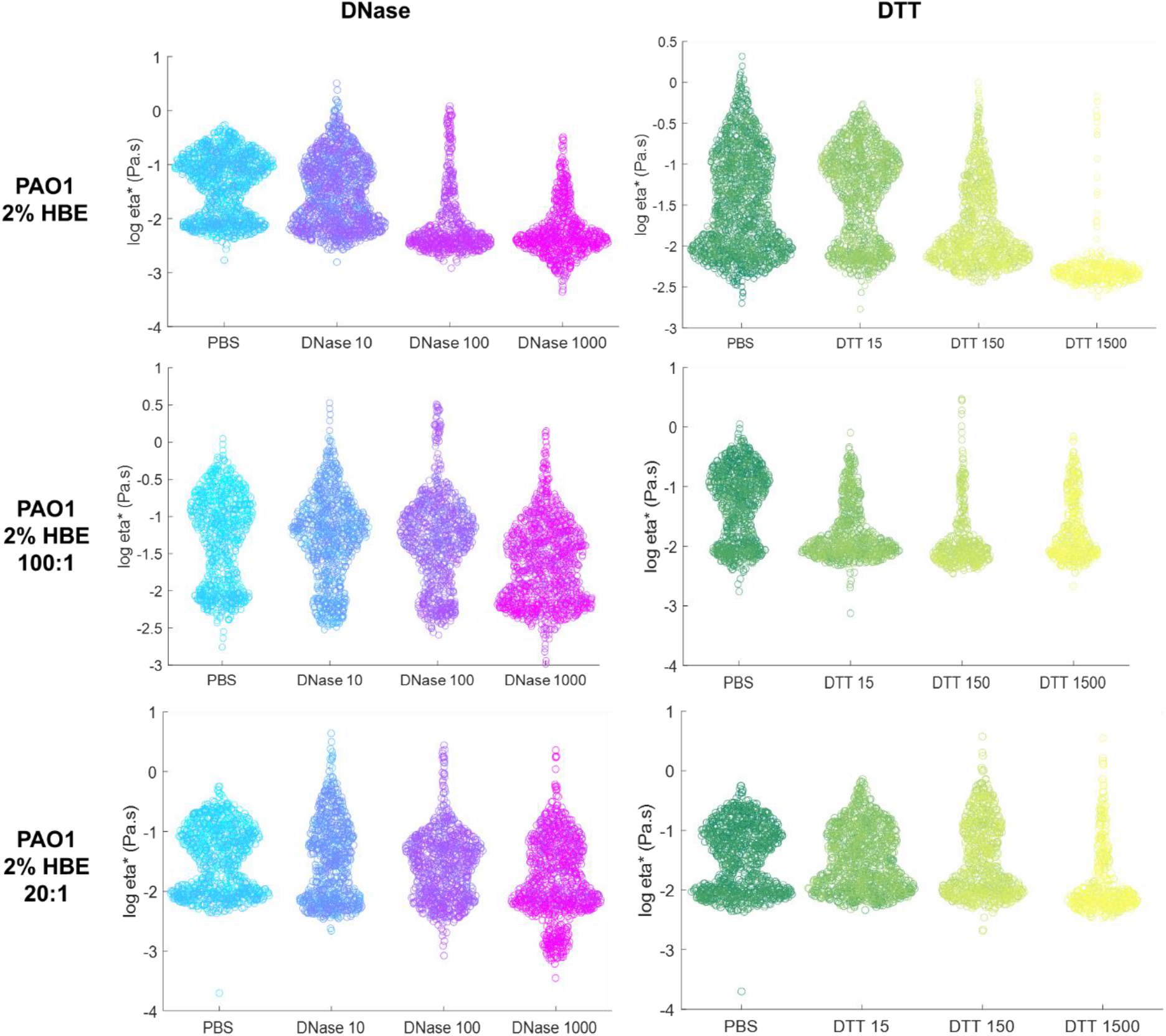
Biofilms grow in 2% HBE microrheology after 24 h exposure to DNase or DTT treatment (μg/mL). Data is presented as a swarm chart of all tracked bead η* in ≥3 separately grown and treated biofilms.

**Figure 5.**
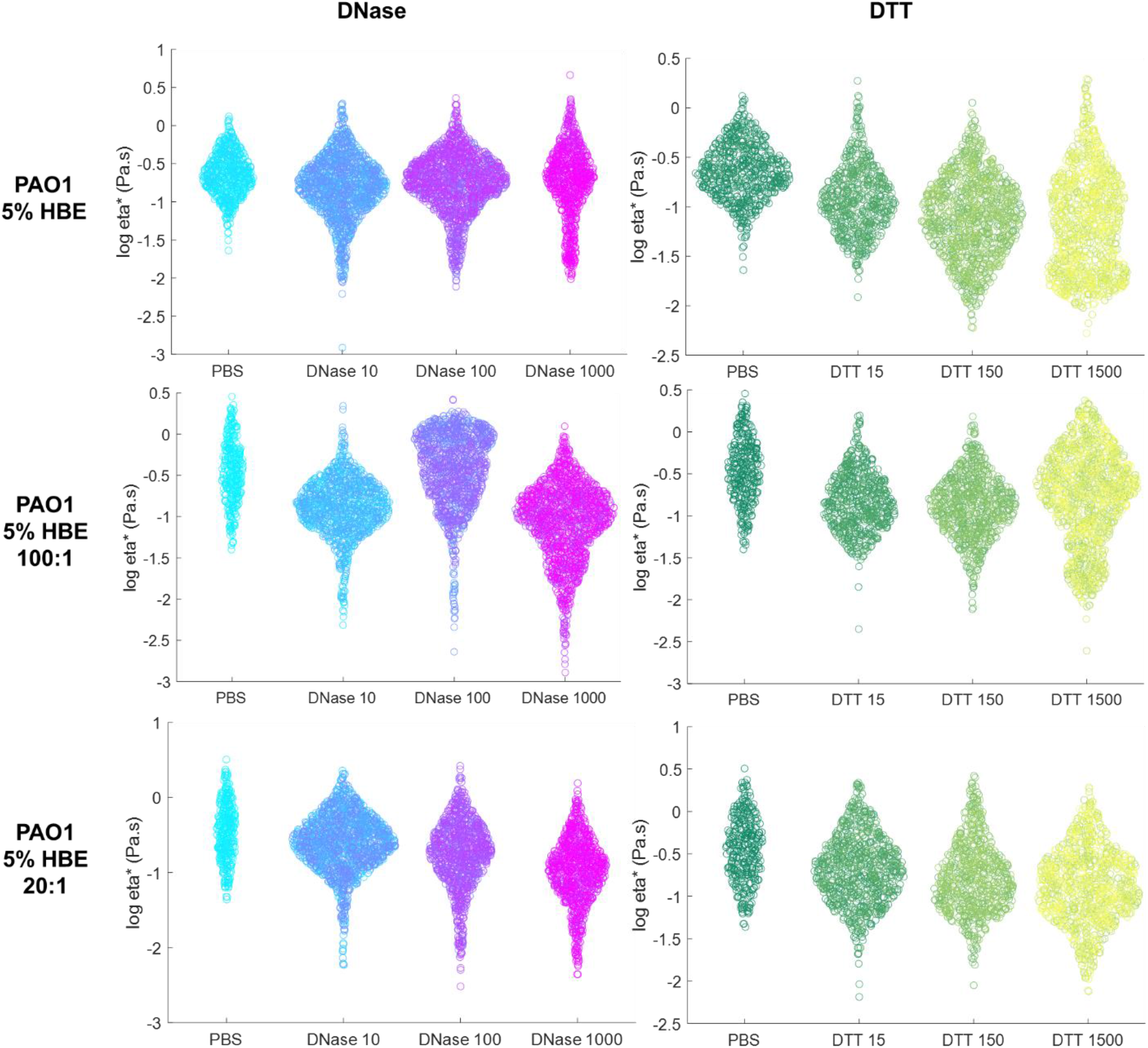
Biofilms grow in 5% HBE microrheology after 24 h exposure to DNase or DTT (μg/mL) treatment. Data is presented as a swarm chart of all tracked bead η* in ≥3 separately grown and treated biofilms.

Indeed, the effects of high dose DNase were apparent in histograms of η* in Figure 6. Untreated 2% biofilms had a distinct bimodal distribution into low and high viscosity components. Treatment with DNase reduced the proportion of signal in the higher η*component, shifted the absolute mean η* value down closer to water, and increased the relative proportion of the low η* component. Biofilms grown in 2% 20:1 also included a component with the viscosity of water after exposure to DNase, indicated by the dashed line. Biofilms grown in 5% mucus types have a single peak with PBS treatment, but upon exposure to DNase, a lower η* component appears in the histogram.

**Figure 6.**
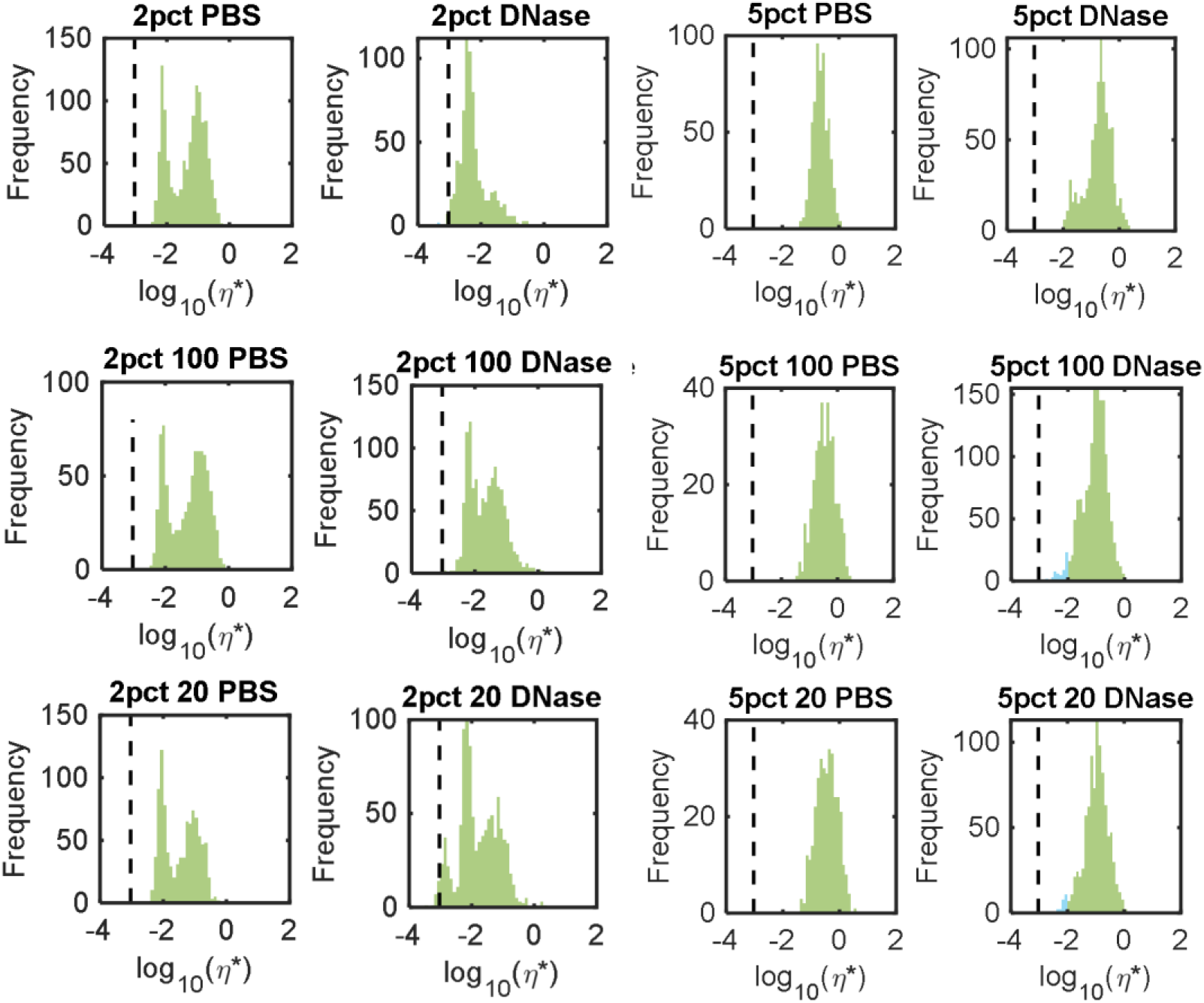
Biofilm microrheology after exposure to PBS or 1000 μg/mL DNase. Data is representative of all tracked bead viscosities in ≥3 separately grown and analyzed biofilms. The concentration of HBE is listed first and 100 or 20 denotes the ratio of mucin to DNA. The dashed line indicates the viscosity of water.

## Discussion

This study represents the first examination into the relationship between mucus concentration/composition and *P. aeruginosa* biofilm mechanical properties. The contributions of mucus concentration and composition to biofilm strength are critical to understand due to the increasing concentration of mucins and DNA with CF pulmonary disease progression. We hypothesized that the viscoelastic properties of *Pseudomonas aeruginosa* biofilms grown in mucus would be dominated by the overall mucus concentration and composition, similar to mucus (4, 16, 17). Indeed, *P. aeruginosa* biofilms grown in nutrient broth (TSB) behaved as a viscoelastic solid (tanδ < 0.5), with G’ greater than G” at 0.64 and 0.11 Pa, respectively (Table 1). Regardless of media composition (i.e., with or without mucins and DNA), all *P. aeruginosa* biofilms studied behaved as viscoelastic solids, likely due to the prevalence of strong chemical and physical interactions in the extracellular matrix between polymers such as DNA and polysaccharides. Consistent with previous reports (27–29), low concentrations of PGM (0.5 and 5 mg/mL) in TSB inhibited robust biofilm formation, as G’ was about half that of those grown in TSB alone. In contrast, environmental DNA alone did not significantly affect biofilm rheology, neither at 0.5 nor 5 mg/mL. However, the combination of PGM and DNA in TSB facilitated robust biofilm formation with macrorheology values 3-4x greater than those grown in TSB alone (Table 1). Thus, the combination of mucins and DNA are associated with mechanically robust biofilms while the individual polymers are not.

Biofilms utilize both endogenously produced and environmentally derived DNA for mechanical stability (18). Treatment with DNase, the DNA degradation enzyme, enzymatically cleaves DNA in the biofilm EPS and alters viscoelastic properties. While the lowest dose of DNase tested by microrheology, 10 μg/mL, had no significant impact on η*, 20 μg/mL increased η* compared to the PBS-treated control. By degrading DNA in the extracellular matrix, chemical bonds are broken, interactions are disrupted, and biofilm components (e.g., cells, biomacromolecules) are released from the interior. Biofilm heterogeneity (κ) increased and EPS rearrangement due to minimal DNA degradation likely contributed to greater η*. Biofilm η* was greater compared to the PBS-treated control but decreased compared to biofilms exposed to 20 μg/mL DNase, suggesting a threshold concentration for uniform biofilm degradation for these biofilms. Biofilm heterogeneity is of significant concern when treating with antibiotics due to the risk of fostering antibiotic resistance when the biofilm is not completely eradicated (3, 33), thus the non-Gaussian parameter of treated biofilms may serve as a quantitative measure of biofilm homogenization. Complete biofilm degradation to the viscosity of water was not achieved with any tested dose of DNase, which indicates that greater concentrations may be necessary or that DNA degradation alone is insufficient to fully degrade biofilms. Combinations of antibiotics and DNase have previously been investigated for superior biofilm eradication (34) and future work will investigate these combinations in mucus-grown biofilms.

The presence of both mucins and DNA together promotes robust biofilm development and targeting DNA in the biofilm matrix allows for partial biofilm degradation. Human bronchial epithelial (HBE) mucus was then used as a growth medium for *P. aeruginosa* biofilms using salmon sperm DNA at physiologically relevant mucin to DNA ratios (17, 24). First, the viscoelastic properties of mucus as a function of concentration and composition were evaluated, and more concentrated mucus was characterized by greater viscoelastic moduli, with 5% being 2-3x greater than 2% (Table 2). The addition of DNA also increased G’, G”, and η*. While the absolute values of 5% mucus η* trended with DNA concentration, both 5% 100:1 and 5% 20:1 mucus were statistically indistinguishable from 5% mucus without DNA on both the macro- and microrheological scales. Mucus physical properties was further resolved with PTMR (Figure 2) where 2% mucus exhibited the bimodal behavior characteristic of healthy mucus (32). The bimodal behavior was mirrored in 2% 100:1 and 2% 20:1, though the curves shifted to greater η*. The prevalence of the lower η* peak (more water-like) was also reduced in mucus with DNA, indicating that 2% mucus with DNA was dominated by the greater η*, or more solid-like, component. Previous work demonstrated that concentrated CF mucus facilitated biofilm formation while 2% mucus did not (5), suggesting the need for a sufficiently robust viscoelastic behavior (i.e., threshold G’, G”, or η* values) in order to allow biofilms to form in mucus.

Indeed, *P. aeruginosa* biofilms grown in 5% were characterized by greater viscoelastic moduli than biofilms grown in 2% by about 4x. Uniquely, biofilms grown in all 2% mucus types exhibited nearly uniform bimodal distribution into lower and higher η* components (Figure 3). However, similar to mucus, the most significant contributor to biofilm rheology was overall mucus concentration (Figures 2 and 3). Significant (p<0.05) increases in macroscopic and microscopic moduli were observed between 2% and 5% mucus and in biofilms grown therein. The addition of DNA was not associated with any significant changes in macroscopic or microscopic moduli compared to mucus or biofilms without DNA. Thus, while DNA is essential to biofilm mechanical stability, the presence of environmental DNA in mucus does not significantly impact the structural integrity of these *P. aeruginosa* biofilms. In fact, only overall mucus concentration was correlated with biofilm macroscopic and microscopic moduli.

Biofilms were exposed to DNase to degrade the DNA in the matrix or to DTT, which is a reducing agent. Both agents degrade biomolecules in the extracellular matrix and disrupt interactions. For simplicity, only biofilm η* will be discussed here. In biofilms grown in 2% mucus ± DNA, DNase at 100 μg/mL consistently reduced macroscopic biofilm η*(Table 4). However, at 1000 μg/mL DNase, the η* of 2% HBE biofilms more than doubled. Biofilms grown in 2% mucus supplemented with DNA were characterized by decreased η* after exposure to 1000 μg/mL DNase (Table 4) at both 100:1 and 20:1 mucin to DNA ratios. The increase in biofilm η* is likely due to cell lysis and the release of cells and stiff cellular debris that results in a greater measured bulk η*, which has previously been described (18). Indeed, PTMR of exposed 2% biofilms demonstrated that 1000 μg/mL DNase eliminated the higher η* component of the biofilm and left a single low η* peak (Figure 4), suggesting that while the local microviscosity of the biofilm decreased, bulk η* may have increased due to the release of cellular debris. In the 2% 100:1 and 2% 20:1 biofilms where the concentration of DNA was greater, DNase degraded DNA in the EPS and decreased bulk biofilm η* at 1000 μg/mL DNase. All biofilms grown in 2% mucus types were similarly affected by 1000 μg/mL DNase on the microrheology scale, with a decrease in the higher η* peak and an increase and/or shift of the lower η* peak (Figure 5). Indeed, biofilms grown in 2% HBE with DNA at both concentrations were degraded by 1000 μg/mL DNase and exhibited smaller macroscopic η*. In contrast, biofilms grown in 2% HBE without DNA treated with 1000 μg/mL DNase exhibited increased macroscopic η*. We therefore posit that a threshold concentration of DNase is required to degrade DNA in the EPS without lysing bacterial cells, whereupon cellular components are released into the bulk and increase macroscopic viscoelasticity. A similar phenomenon was observed in biofilms grown in 5% mucus types. Macrorheological parameters *increased* with low (10 μg/mL) to medium (100 μg/mL) doses of DNase and *decreased* at 1000 μg/mL (Table 4) while PTMR demonstrates a dose-dependent decrease in biofilm η* (Figures 4 and 5).

While DTT physically disrupts *P. aeruginosa* biofilms, no consistent trend across mucus types was observed with DTT treatment, which reduces chemical bonds including disulfide bonds. In 2% mucus biofilms, macroscopic moduli increased with DTT concentration (Table 4). The opposite effect was observed in 2% 100:1 biofilms. In 2% 20:1 biofilms, macroscopic η* dramatically increased with the lowest tested dose of DTT and then decreased below the value for the PBS-treated control with greater doses of DTT. The lack of an observable trend in biofilm macrorheology may be due in part to the toxicity of DTT, which generates reactive oxygen species (ROS) capable of damaging DNA (35). Thus, any changes in biofilm rheology due to the reduction of chemical bonds (e.g., mucin disulfide bonds) in the extracellular matrix may be hidden by changes in rheology due to cell death and ROS reactions. Further, bulk rheology may be affected by cell lysis and the release of exoenzymes, swellable polysaccharides, proteins, and lipids as the biofilm is degraded (13, 18). Biofilm disruption with antibacterial agent treatment has previously been associated with increases in macroscopic moduli (18) thus PTMR was used to evaluate changes in biofilm microrheology with DTT treatment. Microrheology of DTT-treated biofilms grown in 2% mucus types showed the presence of a tall and narrow peak at a η* near water (Figure 4) suggesting that the physical biofilm had been sufficiently degraded for the 1 μm probe to measure a predominantly low η* fluid. In biofilms grown in 5% mucus types, DTT treatment was associated with an increase in biofilm heterogeneity and downward shift in η*, suggestive of biofilm degradation. As macrorheology is dominated by the bulk rheological response, microrheology is consistently superior for investigating the biophysical effects of DNase and DTT treatment on *P. aeruginosa* biofilms. Together these data suggest that biofilm disruption alone is insufficient to consistently reduce biofilm η* to that of water, which would, in theory, contribute to improved clearance. Further work is needed to characterize the relationship between biofilm viscoelasticity and mucociliary clearance. Also, less cytotoxic reducing agents must be evaluated to better understand the impact of disulfide bond reduction between mucins in mucus-grown biofilms.

In summary, the predominant contributor to biofilm rheology was overall mucus concentration. The presence of DNA in mucus is associated with greater mucus viscoelasticity but not necessarily increased biofilm mechanical strength. Decreasing the concentration of airway mucus in CF patients may help prevent the establishment of chronic respiratory infections. Regardless of the DNA concentration in mucus, *P. aeruginosa* biofilm microrheology was reduced by DNase and DTT treatments, though bulk η* increased with biofilm disruption. Thus, treatments that target both DNA and EPS interactions may be superior antibiofilm agents for CF respiratory infection, though polymer degradation alone may be insufficient for biofilm eradication. Biofilm mechanical strength has previously been associated with susceptibility to antibiotics (2, 11, 18). As both DNase and DTT are capable of physically degrading *P. aeruginosa* biofilms, their concomitant use with an antibacterial agent may facilitate superior biofilm eradication. Additionally, physical degradation of the biofilm may assist the host immune response in clearing the biofilm via phagocytosis or other clearance mechanisms. To prevent chronic infection, the modulation of mucus concentration will be critical. However, once chronic infection is established, physical biofilm disruption in combination with mucus hydration and mucolysis will promote eradication and clearance. Future work will investigate the combination of biofilm degradation and antibacterial action using antibacterial agents that are effective regardless of bacteria metabolism and evaluate mucociliary transport rates with treatment.

## Methods

### Materials

All materials were used as received unless otherwise specified. Fluorescent 1 μm carboxylated latex beads (FluoSpheres™, Invitrogen), tryptic soy broth (TSB), tryptic soy agar (TSA), porcine gastric mucins (PGM), salmon sperm DNA, DNase, and dithiothreitol (DTT) were purchased from Millipore Sigma. BacLight Green and propidium iodide (PI) were purchased from ThermoFisher. The *P. aeruginosa* strain PAO1 was obtained from the American Type Culture Collection (Manassas, VA), ATCC #15692.

### Biofilm growth in nutrient broth and treatment

Tryptic soy broth (TSB) was inoculated with *P. aeruginosa* strain PAO1 frozen stock and cultured overnight. The overnight culture was resuspended in fresh TSB and cultured at 37 °C to an OD_600_ of 0.25. Bacteria were further diluted in fresh TSB to a final concentration of 10^6^ CFU/mL and 150 μL of this solution was added to the wells of a 96-well plate. The well plate was sealed with parafilm and incubated with shaking at 37 °C for 3 d until a viscous macrocolony formed as previously described (36). Tryptic soy broth was also supplemented with porcine gastric mucins (PGM, 5 mg/mL) and salmon sperm DNA (4 mg/mL), either alone or in combination to match concentrations used in artificial sputum media (ASM) (25). The biofilm (50 μL) was removed from growth media and added to a microcentrifuge tube prior to treatment or analysis using a positive pressure pipette as previously described (18). Dissolved DNase in PBS was added in a volume of 5 μL or less into the center of the biofilm to final concentrations ranging from 10 to 1000 μg/mL. Microcentrifuge tubes were incubated with shaking at 37 °C for 24 h prior to analysis.

### Mucus preparation and biofilm growth

Human airway epithelial cells were obtained by the Marsico Lung Institute Tissue Procurement and Cell Culture Facility and cultured at the air-liquid interface as previously described (16). Mucus washings were collected, pooled, and dialyzed. Total % solids was determined via dry weight calculations, and mucin content was quantified with refractometry. Mucus (100 μL) was added to a 96 well-plate and inoculated with *P. aeruginosa* to a final concentration of 10^6^ CFU/mL. For particle tracking microrheology (PTMR) experiments, 2 μL of carboxylated latex beads at a 1:10 dilution from stock was added (18). Adjacent wells were filled with 200 μL distilled water to prevent evaporation, and the lid was sealed to the plate with parafilm. The sealed well plate was incubated with shaking at 37 °C for 24 h until a visible, viscous microcolony formed at the air-liquid interface as previously described.(5, 9, 18, 37) The biofilm (50 μL) was removed from growth media and added to a microcentrifuge tube prior to treatment or analysis. Biofilms were treated with DNase or dithiothreitol (DTT) in concentrations ranging from 10 to 1500 μg/mL.

### Rheology

Macrorheology was performed using a 20 mm 1 ° cone and plate rheometer and frequency sweeps as previously described (18). Particle tracking microrheology was performed with 1 μm carboxylated latex beads and video microscopy as previously described (18, 32). All measurements were repeated in biological duplicate with five technical replicates, and data is reported as the mean ± standard deviation. All values of frequency-dependent parameters are given as a single value from a frequency sweep assay with the value reported corresponding to the measured response at 1 Hz.

### Statistics

Each measurement was repeated ≥3 times per sample. Mucus samples were separately prepared and analyzed. Biofilm samples were grown on different days using a fresh overnight solution. Data is presented as the mean and standard deviation. Statistical significance was determined using a two-tailed Student’s *t*-test.

## Acknowledgments

Dr. Kaitlyn R. Rouillard is funded through the T32 program, Multidisciplinary Research Training in Pulmonary Diseases, T32HL007106, contact PI Dr. Claire Doerschuk. This research was supported by the Cystic Fibrosis Foundation (HILL20Y2-OUT, HILL19G0, BOUCHE19R0).

